# Contextual Scaling of Representational Uncertainty for Size and Number

**DOI:** 10.1101/2025.10.31.685954

**Authors:** Yosuke Sakamoto, Masamichi J. Hayashi

**Author notes:** Correspondence concerning this article should be addressed to Masamichi J. Hayashi, Center for Information and Neural Networks (CiNet),1-4 Yamadaoka, Suita, Osaka 565-0871, Japan. **Author Note** The authors declare no competing interest.

## Abstract

Accurate discrimination of magnitudes, such as object size and quantity, is crucial for effective action selection and decision-making. A fundamental principle in magnitude discrimination is that discriminability is determined by the ratio of the stimulus values being compared, as formalized by Weber-Fechner’s law. Here, we show evidence that the discriminability of numerosity is not as robust as previously thought; rather, it is influenced by task-irrelevant environmental statistics. We measured discrimination thresholds in 24 participants performing a numerosity discrimination task in which pairs of dot arrays were presented either simultaneously or sequentially. Stimulus values were sampled from either narrow or wide ranges. Our findings revealed that, although the magnitude pairs were identical, discrimination performance was significantly better (i.e., lower threshold) when stimulus values were sampled from a narrow-range compared to a wide-range. Strikingly, this improvement was observed only when the stimuli were presented sequentially. A similar pattern emerged in a size discrimination task with analogous manipulations. A control experiment ruled out potential confounding variables related to stimulus variety, emphasizing the importance of the stimulus range in gaining this benefit. Together, these findings suggest that perceptual variability is flexibly scaled according to context, particularly when the representations are less reliable. Our study highlights the adaptive nature of stimulus encoding strategies, whereby the brain minimizes representational uncertainty by dynamically leveraging contextual information.

---

Processing size and number is crucial for guiding behaviors such as foraging (Bonn & Cantlon, 2012; Tsouli et al., 2022) and economic decision-making (Barretto-García et al., 2023). A hallmark of magnitude processing, including size and numerosity, is that the discriminability between two magnitudes is determined by their distance in logarithmic space (Ditz & Nieder, 2016; Nieder & Miller, 2003; Piazza et al., 2004; Tsouli et al., 2022). This characteristic is formalized by Weber-Fechner’s law, which posits that the variability of internal representations increases in proportion to magnitude (Tsouli et al., 2022). Notably, this principle is preserved across a wide variety of species, including non-human primates (Nieder & Miller, 2003) and birds (Ditz & Nieder, 2016), suggesting an evolutionarily robust basis of magnitude encoding.

At the neural level, activity in the human frontoparietal regions, which are thought to support magnitude processing across different dimensions (Walsh, 2003), is tuned to specific preferred magnitudes (Harvey et al., 2015; Hayashi et al., 2015; Piazza et al., 2004; Tudusciuc & Nieder, 2007). For example, numerosity-tuned neural populations exhibit bell-shaped tuning curves characterized by a preferred magnitude (i.e., the peak of the curve) and its associated variability (i.e., the tuning width; Piazza et al., 2004; Tudusciuc & Nieder, 2007). Crucially, this tuning width increases with preferred magnitude, mirroring the behavioral signature of Weber-Fechner’s law (Ditz & Nieder, 2016; Nieder & Miller, 2003; Piazza et al., 2004). This correlation suggests a close relationship between behavioral performance and the underlying neural mechanisms.

It is well established that numerosity representations are highly adaptive to context (Kido et al., 2025; Namdar et al., 2016; Piazza et al., 2004; Tsouli et al., 2021). For instance, perceived numerosity is biased after repeated exposure to a specific numerosity (i.e., adaptation; Burr & Ross, 2008) and is also attracted toward previously presented numerosity through serial dependence (Cicchini et al., 2014; Fornaciai & Park, 2018). Over a broader timescale, numerical estimates tend to be biased toward the center of the experienced numerosity range (i.e., central tendency; Anobile et al., 2012, 2019). Although the neural bases of the perceived numerosity remain unclear, such contextual modulation is likely to modulate responses in numerosity-selective regions (Castaldi et al., 2016). Moreover, a recent neuroimaging study demonstrated a context-dependent coding strategy for numerical magnitude (Kido et al., 2025). Specifically, the preferred numerosities of neurons in the frontoparietal cortex may be scaled according to the numerical range of the presented stimuli. These findings indicate that magnitude representations are not fixed, but rather adapt to the statistical structure of the environment, such as stimulus distributions.

While these previous studies have primarily focused on context-dependent biases in numerosity representations, the influence of stimulus range on representational variability remains unclear. Given that variability is linked to perceptual discriminability, any context-dependent modulation of variability should be reflected in discrimination performance. Specifically, when observers are required to encode a narrower range of magnitudes, they may reduce representational variability to increase precision. Here, through a series of pre-registered psychophysical experiments, we demonstrate that the variability of numerical magnitude representations flexibly scales with contextual range, and that this trend is evident not only in numerosity but also in size discrimination. This systematic departure from the constant relative precision predicted by Weber-Fechner’s law reveals a domain-general mechanism of adaptive encoding that operates as a function of contextual range.

## Materials and Methods

Prior to data collection, we pre-registered the experimental plans, including sample size, exclusion criteria, procedure, and analysis described below at *AsPredicted* (https://aspredicted.org/w5xj-tkgn.pdf for Experiment 1 and https://aspredicted.org/vkgb-3grf.pdf for Experiment 2). The study was approved by the institutional ethics and safety committees of the National Institute of Information and Communications Technology.

### Participants

For both Experiments 1 and 2, we aimed to collect data from 24 healthy adult volunteers who passed our exclusion criteria (see below). We determined this sample size based on a priori power analysis (G*Power; Faul et al., 2009) using the smallest effect size observed in a pilot study (η^2^ = 0.09), with a significance level of 0.05 and statistical power of 80 %. We recruited participants using the same procedure across experiments. For the first round, 24 participants were recruited and participated in the experiment. After applying the pre-determined exclusion criteria, we recruited the excluded number of participants and then reapplied the same criteria. This process continued until the pre-determined sample size was achieved.

We finalized our results based on analyses of data from 24 participants in Experiment 1 (19 males, 5 females, mean age: 22.55, *SD* = 1.31) and another group of 24 participants in Experiment 2 (16 males, 8 females, mean age: 22.96, *SD* = 2.24). Participants completed two sessions in Experiment 1 and one in Experiment 2, and were paid 3,000 yen per session.

All participants were right-handed and naïve to the purpose of the experiment. They reported no history of neurological or psychiatric disorders and had correct or corrected-to-normal visual acuity. We obtained written informed consent from all participants.

### Exclusion Criteria

We excluded participants based on the following criteria:

a. Participants who do not complete all the tasks.
b. Participants who do not make a response in more than 5 % of entire trials in any experimental conditions and tasks.
c. Participants whose psychometric function in any experimental conditions and tasks fails to converge.
d. Participants whose slope of psychometric function deviates from the across-participant mean ± 3 standard deviation (SD) in any experimental conditions and tasks.
e. Participants whose point of subjective equality (PSE) of psychometric function deviates from the across-participant mean ± 3SD in any experimental conditions and tasks.
f. Participants whose task accuracy in any experimental conditions and tasks is lower than 60% of all valid trials.

### Task and Stimuli

Participants performed a numerosity discrimination task and a size discrimination task (**Figure 1A**). Each trial began with a white fixation cross (0.5° × 0.5°, 500 ms duration) presented at the center of the screen. Subsequently, a pair of dot arrays for the numerosity task or disks for the size task (**Figure 1B**), consisting of a standard and a comparison stimulus, was presented either simultaneously or sequentially. In the simultaneous condition, the stimulus pair was presented for 300 ms in the right and left visual fields; the position of the standard stimulus, right or left, was balanced within each block. In the sequential condition, the two stimuli were presented sequentially, one on each side of the visual field, for 300 ms each, with a 1,000-ms inter-stimulus-interval. The order of the standard and comparison stimuli and the position of the first-presented stimulus were balanced within each block. After stimulus presentation, the fixation cross turned red, initiating a response period during which participants indicated which of the two stimuli contained the larger number of dots in the numerosity task or was larger in size in the size task by pressing one of two response keys. Participants were instructed to maintain their gaze at the fixation cross throughout each experimental block and to respond as quickly as possible. The next trial was initiated by the participant’s response or after 2,500 ms had elapsed without a response.

**Figure 1.**
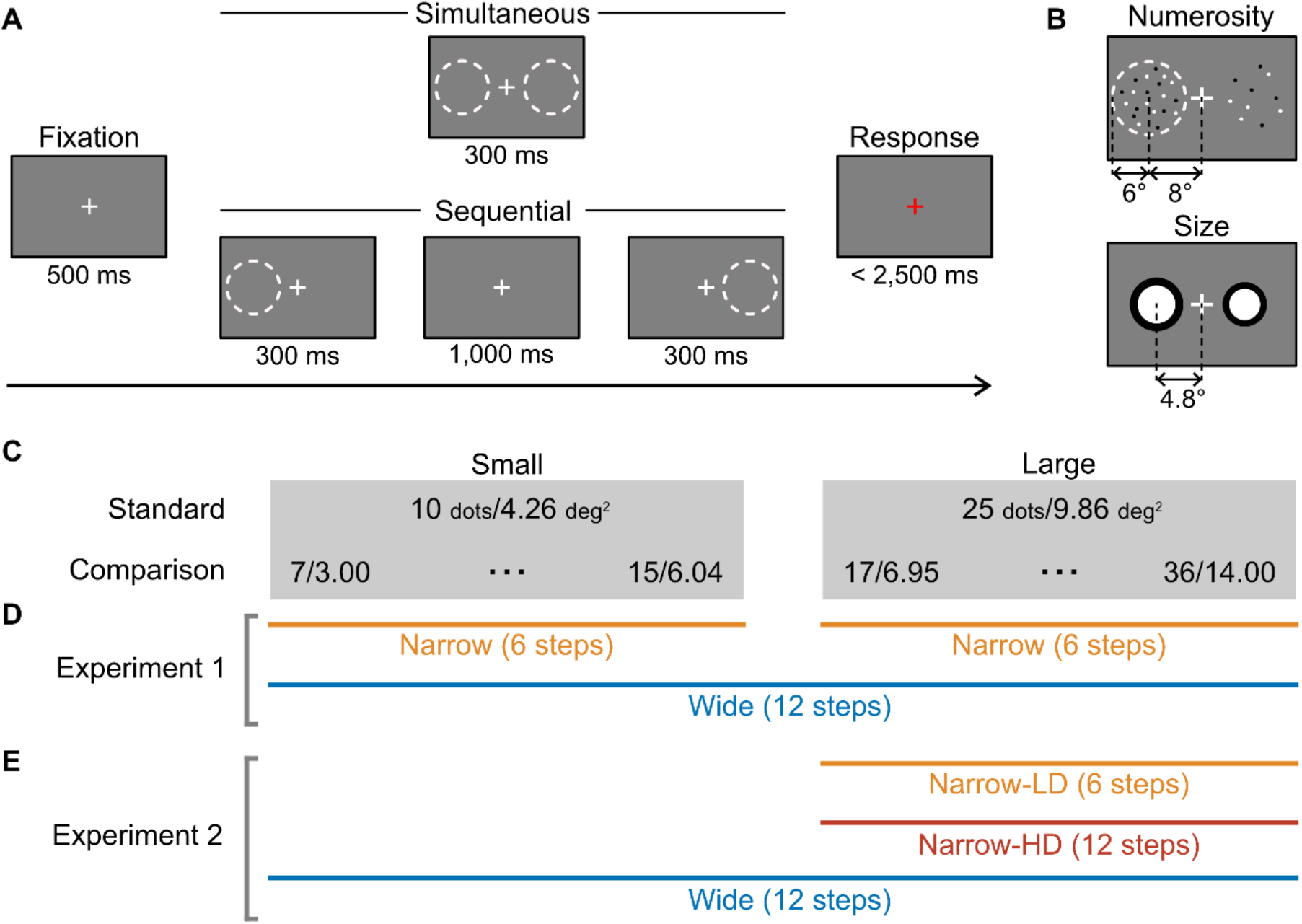
Experimental design. (A) Stimulus sequence of a trial. (B) Example stimuli for the numerosity (top) and size (bottom) task. (C) Stimulus magnitude parameters. Stimulus values for the numerosity task (number of dots) and size task (area in deg^2^) are shown for small and large magnitude levels. (D, E) Schematic illustration of the stimulus range conditions used in Experiment 1 (D) and Experiment 2 (E).

In Experiment 1, participants performed the numerosity and size discrimination tasks at two standard magnitude levels: small, corresponding to 10 dots or 4.26 deg^2^, and large, corresponding to 25 dots or 9.86 deg^2^ (**Figure 1C**). The comparison stimulus values were pseudo-randomly selected on each trial from six logarithmically spaced steps: 7-15 dots or 3.00-6.04 deg^2^ for the small level, and 17-36 dots or 3.00-6.04 deg^2^ for large level (**Figure 1C** and **1D**). For numerosity stimuli, these values were rounded to the nearest integer. The stimulus pairs were presented in two different contextual ranges: narrow-range and wide-range conditions. In the narrow-range condition, stimulus pairs from only one of the two magnitude levels were presented throughout an experimental block. In contrast, in the wide-range condition, stimulus pairs from the two magnitude levels were randomly interleaved on a trial-by-trial basis. The number of trials were equivalent for all stimulus pairs, resulting in a uniform distribution.

Experiment 2 was identical to Experiment 1 except for the following modifications. First, stimuli were always presented sequentially. Second, in addition to the wide- and narrow-range conditions, the latter hereafter referred to as the narrow-low-density condition, or narrow-LD, we introduced an additional condition in which the stimulus range was identical to that of the narrow-LD condition at the large magnitude level, but the number of comparison-stimulus steps was identical to that in the wide-range condition; that is, the narrow-range was divided into 12 rather than 6 steps (**Figure 1E**). We refer to this new condition as the narrow-high-density condition, or narrow-HD. Thus, in Experiment 2, each task consisted of three experimental blocks: wide-range, narrow-LD, and narrow-HD.

The dot-array stimuli used for the numerosity task consisted of white and black dots, each 0.2° in diameter in visual angle, and were presented within an invisible circle with a diameter of 6°, centered 8° to the right or left of the screen center (**Figure 1B**). The positions of individual dots within the invisible circles were randomized across trials, while maintaining a minimum inter-dot distance of 1° to prevent overlap (Park, 2022). We chose these stimulus parameters so that the number of dots fell within the numerosity range, which is thought to rely on mechanisms distinct from those underlying subitizing and texture-density-like processing (Burr et al., 2018). The numbers of white and black dots were equal when an even number of dots was presented. When an odd number of dots was presented, one of the two colors had one more dot than the other, and the color with the extra dot was counterbalanced across trials. The disk stimuli used for the size task consisted of two concentric circles and were presented on the left and right sides of the screen, centered 4.8° to the left and right of the screen center. The area of the inner white disk was equal to that of the outer black ring.

All visual stimuli were presented on a grey background using a 24-inch gamma-corrected LCD monitor (BenQ E2420HD; 60 Hz refresh rate, 1920 × 1080 resolution) at a viewing distance of 50 cm. We validated the timing of stimulus presentation using an oscilloscope. PsychoPy 3 v2024.2.3 (Peirce et al., 2019), implemented in Python 3.10, was used to generate and present the visual stimuli.

### Procedure

#### Experiment 1

The experiment was conducted across two sessions, separated by the presentation paradigm (simultaneous or sequential), with the order counterbalanced across participants. In each session, participants completed both numerosity and size discrimination tasks. Each task contained four experimental blocks consisting of two consecutive wide-range condition blocks and two narrow-range condition blocks, each corresponding to the small and large levels. The task and contextual range orders were fixed for a given participant across sessions but counterbalanced across participants.

Participants completed 24 practice trials prior to each experimental block, except for the second wide-range condition block. Feedback for response correctness was provided only during the practice trials, signed by the colored square (green: correct; red: incorrect) presented for 500 ms at the end of each trial. Each session consisted of 768 trials (6 comparison stimuli × 32 repetitions × 4 blocks). Experiments lasted up to 120 minutes per session.

#### Experiment 2

Experimental procedure was identical to Experiment 1, except for the number of experimental blocks (four in Experiment 1 and three in Experiment 2). Comparison stimuli were presented with 16 repetitions for each block, resulting in 192 trials for wide-range (6 comparison stimuli × 16 repetitions × 2 magnitude levels) and narrow-HD (12 comparison × 16 repetition), and 96 trials for narrow-LD (6 comparison × 16 repetition) conditions.

### Analysis

We conducted all analyses using R (version 4.3.3). To quantify discrimination performance, we fitted psychometric functions to individual data using the *lme4* package (Bates et al., 2015) and then estimated the Just Noticeable Difference (JND) from the fitted psychometric curves. We performed three-way repeated measures ANOVAs (simultaneous/sequential paradigm × wide-/narrow-range × small/large level) for Experiment 1 and one-way repeated measures ANOVAs (wide-range/narrow-LD/narrow-HD condition) for Experiment 2, separately for numerosity and size discrimination tasks. All statistical tests were performed at a significance level of 0.05, and results from the post-hoc two-tailed *t*-tests were corrected using Shaffer’s method for multiple comparisons.

## Results

In Experiment 1, we tested whether the discrimination performance improves when stimulus values are selected from a narrower magnitude range. We found that the JND for numerosity discrimination was modulated by the interaction between the experimental paradigm and stimulus range (*F*(1, 23) = 8.92, *p* = .0066, *Cohen’s f* = 0.079). Specifically, participants exhibited a smaller JND (indicating better discriminability) in the narrow-range condition when stimuli were presented sequentially (*F*(1, 23) = 11.53, *p* = .0025, *f* = 0.12). However, no such improvement was observed when stimuli were presented simultaneously (*F*(1, 23) = 0.67, *p* = .42, *f* = 0.032; **Figure 2A**). The same trend was observed in the JND for the size discrimination task with a significant interaction (*F*(1, 23) = 18.70, *p* < .001, *f* = 0.11), showing better discrimination performance in the narrow-range condition under sequential presentation (*F*(1, 23) = 25.13, *p* < .001, *f* = 0.21), and no improvement under simultaneous presentation (*F*(1, 23) = 0.04, *p* = .84, *f* = 0.01; **Figure 2C**). In summary, the variability of magnitude representations for both number and size was flexibly scaled by context, but only when magnitude information needed to be temporarily maintained. These results suggest that the human brain adjusts representational uncertainty according to contextual factors; when stimuli are less reliable due to temporary maintenance, the brain adjusts the variability of magnitude representation accordingly.

**Figure 2.**
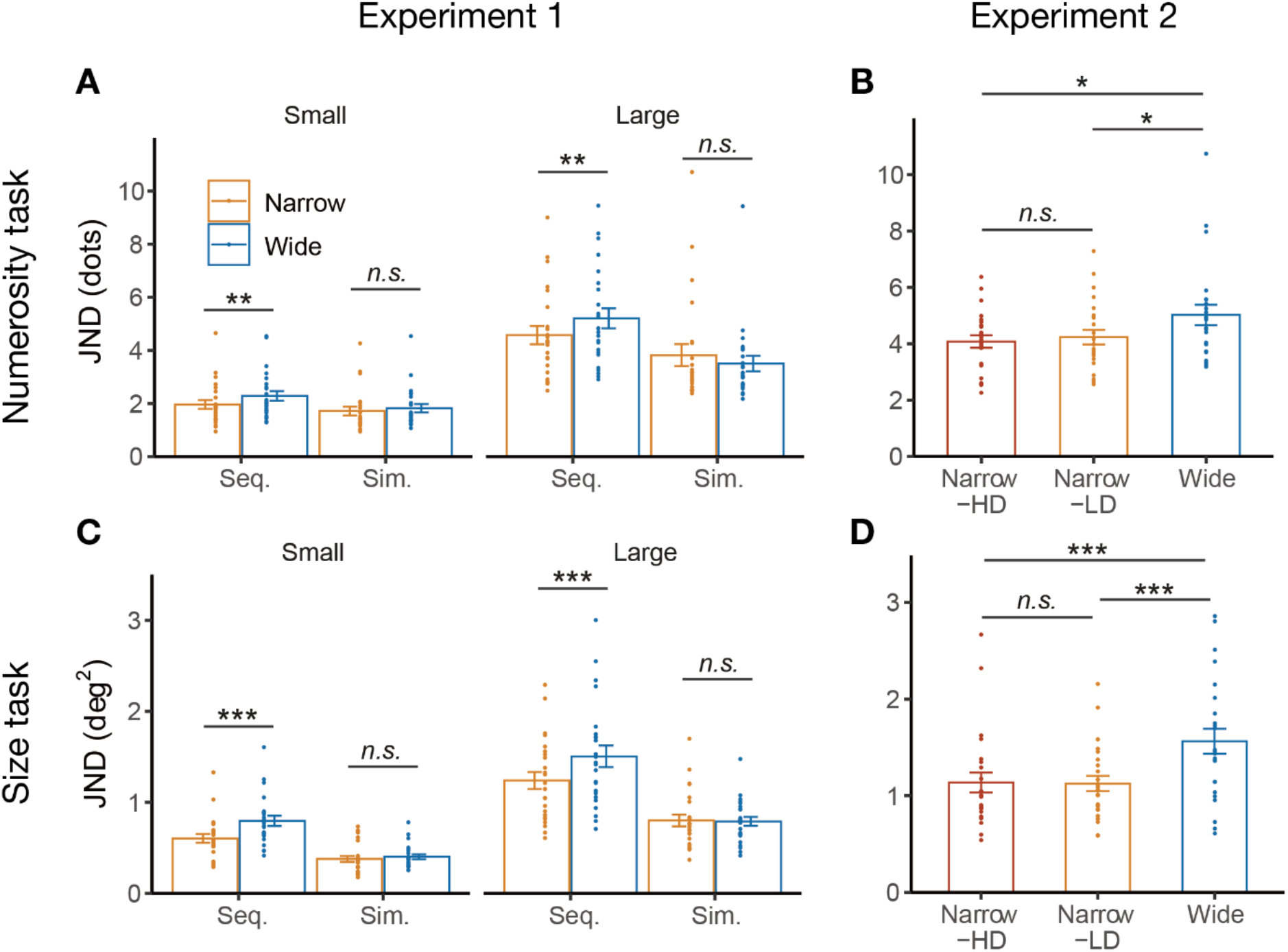
Results of Experiment 1 and 2. (A, C) Mean JNDs across participants for the numerosity (A) and size (C) tasks in Experiment 1. Seq. and Sim. denote the sequential (2IFC) and simultaneous (2AFC) paradigms, respectively. (B, D) Mean JNDs for the large magnitude level across participants in Experiment 2 for the numerosity (B) and size (D) tasks. Colored dots denote individual data (n = 24). Error bars indicate SEM. *p < .05, **p < .01, ***p < .001.

An alternative explanation for the improved discrimination performance in the narrow-range condition could be the smaller number of stimulus varieties compared to the wide-range condition (i.e., six versus twelve values), rather than a difference in the stimulus range. Therefore, we conducted Experiment 2 in which the stimulus range and the stimulus varieties within ranges are carefully controlled (**Figure 1D**). We found that the variety of stimulus values was not relevant to the modulation of thresholds observed in Experiment 1; rather, the range played a significant role (main effect of one-way repeated measures ANOVA, *F*(2, 46) = 5.17, *p* =.0095, *f* = 0.30 for numerosity discrimination; *F*(2, 46) = 15.73, *p* < .001, *f* = 0.40 for size discrimination). We replicated the findings of a smaller JND in the narrow-LD condition compared to the wide-range condition (*t*(23) = 2.20, adjusted *p* = .038, *Cohens’ d* = 0.45 for numerosity task; *t*(23) = 4.52, adjusted *p* <.001, *d* = 0.92 for size task). Importantly, no significant difference was found between narrow-LD and narrow-HD (*t*(23) = 0.69, adjusted *p* = .50, *d* = 0.14 for numerosity task; *t*(23) = 0.20, adjusted *p* = 0.84, *d* = 0.04 for size task). Most critically, the JND in the narrow-HD condition was significantly smaller than in the wide-range condition (*t*(23) = 2.75, adjusted *p* =.035, *d* = 0.56 for numerosity task; *t*(23) = 3.97, adjusted *p* < .001, *d* = 0.81 for size task). These results indicate that the contextual scaling of variability is driven by the range of magnitudes themselves rather than the distinct number of stimulus values to be encoded.

## Discussion

In this study, we found that discrimination performance for both size and numerosity improved when stimulus values were drawn from a narrow range. This effect was especially pronounced when stimuli were presented sequentially, requiring temporary maintenance in WM. Together, these findings suggest that when incoming magnitude information is variable and noisy, the human brain can actively reduce representational variability by exploiting contextual information.

Our findings extend previous works on context-dependent magnitude encoding, such as central tendency effects, which have largely relied on reproduction paradigms, in three important respects. First, whereas earlier studies primarily focused on biases in perceived spatial magnitude (Petzschner & Glasauer, 2011; Xiang et al., 2021), our results show that the discriminability of stimulus magnitudes is also influenced by context, which we manipulated through the range of the stimulus distribution in a discrimination task. Second, by introducing WM demands through two distinct presentation paradigms, we provide evidence that WM maintenance plays a critical role in enabling contextual information to enhance discriminability. Finally, by showing similar results across two spatial magnitude domains, namely size and numerosity, we suggest that modulation of a shared magnitude representations, as proposed by A Theory Of Magnitude (Bueti & Walsh, 2009; Walsh, 2003), may underly the context-dependent improvements in discriminability reported here. Taken together, these findings complement prior studies of context-dependent magnitude encoding by showing that contextual factors can flexibly modulate perceptual precision, particularly when WM is engaged.

These findings can be interpreted within a Bayesian framework (Jazayeri & Shadlen, 2010; Miyazaki et al., 2006; Petzschner et al., 2015; Petzschner & Glasauer, 2011; Roach et al., 2017). In this framework, a narrower stimulus range gives rise to a sharper prior distribution, which in turn produces a stronger contraction of posterior estimates toward the prior mean. Such contractions have been shown to be especially pronounced when the variance of the likelihood function is larger, that is, when sensory information is less reliable (Miyazaki et al., 2005). A similar enhancement can occur when information must be temporarily retained in visual WM as reflected in contraction bias (Tal-Perry & Yuval-Greenberg, 2022). We therefore suggest that these two factors may jointly account for the improved discrimination performance observed in the narrow-range condition in our study, particularly when one of the stimulus representations became less reliable because of temporal maintenance. More specifically, when stimuli were presented sequentially, the presentation of the first stimulus may have been more strongly shaped by prior expectations, leading to greater perceptual precision in the narrow-range than in the wide-range condition. By contrast, when stimuli were presented simultaneously, both stimuli were likely influenced by the magnitude range to a similar degree, thereby attenuating range-dependent differences. We therefore propose that improved behavioral performance arose from the combination of a narrower prior and an asymmetry in the variance of the likelihood functions for the two stimuli.

Although the neural mechanisms underlying this scaling of variability remain an open question, one possibility is that the behavioral effect reflects a sharpening of the tuning of numerosity- and size-selective neural populations by contextual range information, thereby enhancing behavioral precision. This possibility is supported by previous studies showing that prior expectations and adaptation can sharpen sensory population tuning (Kohn & Movshon, 2004; Krekelberg et al., 2006; Park et al., 2023), and that such sharpening is associated with enhanced behavioral precision (Kok et al., 2012; Krekelberg et al., 2006; Park et al., 2023).

Given evidence that neurons in the frontoparietal cortex may encode context-dependent, relative numerical magnitudes (Kido et al., 2025), we speculate that the contextual scaling of variability may similarly arise from context-dependent sharpening of relative magnitude representations. More broadly, this context sensitivity underscores the flexibility of the neurocomputational mechanisms supporting magnitude processing.

## Acknowledgments

We thank Dr. Motoaki Uchimura for his valuable input for this work. This work was supported by the Japan Science and Technology Agency (FOREST JPMJFR232X to M.J.H.) and the Japan Society for the Promotion of Science (Grant-in-Aid for Scientific Research JP22H01110 to M.J.H., Grant-in-Aid for Scientific Research on Innovative Areas JP21H00315 to M.J.H.).

## Declarations

### Ethics approval

This study was approved by the institutional ethics and safety committees of the National Institute of Information and Communications Technology.

### Consent to participate

All participants provided written informed consent.

### Consent for publication

Not applicable.

### Availability of data and materials

Data for the experiments are available upon request.

### Code availability

The scripts for the analyses are available upon requenst.

### Authors’ contributions

**Yosuke Sakamoto**: Conceptualization, Methodology, Software, Validation, Formal analysis, Investigation, Data curation, Writing - original draft, Writing - review & editing, Visualization. **Masamichi J Hayashi**: Conceptualization, Methodology, Validation, Resources, Writing - review & editing, Supervision, Project administration, Funding acquisition.

